# A comparison of mtDNA deletion mutant proliferation mechanisms

**DOI:** 10.1101/2022.05.03.489489

**Authors:** Alan G Holt, Adrian M Davies

## Abstract

In this paper we use simulation methods to investigate the proliferation of deletion mutations of mitochondrial DNA in neurons. We simulate three mtDNA proliferation mechanisms, namely, random drift, replicative advantage and vicious cycle. For each mechanism, we investigated the effect mutation rates have on neuron loss within a human host. We also compare heteroplasmy of each mechanism at mutation rates that yield the levels neuron loss that would be associated with dementia. Both random drift and vicious cycle predicted high levels of heteroplasmy, while replicative advantage showed a small number of dominant clones with a low background of heteroplasmy.

## 1 Introduction

Modest decline of cognitive function with advancing age is accepted as a normal part of the aging process [33]. However, current research indicates that, with no underlying pathology, neuron loss is small [43]. In contrast to healthy aging, dementia is associated with substantial loss of neurons [4, 14, 17, 19, 40].

A key risk factor for neurodegenerative diseases is age [22, 45]. Indeed, the prevalence of dementia increases dramatically past the age of 65 [38]. While there are likely to be numerous factors that influence the onset of dementia, mitochondrial dysfunction is believed to be the cause of various neurodegenerative diseases [41, 42] due to the accumulation of mitochondrial DNA deletion mutations (mtDNA_*del*_) [3, 10, 26, 27, 35].

A primary function of mitochondria is the generation of ATP; Mitochodrial DNA (mtDNA) codes for several necessary components of the electron transport chain. Significant numbers of mtDNA_*del*_ are likely to impair mitochondrial function and cause neuron loss mediated by an energy crisis [44].

mtDNA within neurons have a relatively short half-life which is in the order of 30 days [18, 23]. Neurons, on the other hand, are post-mitotic and must have a lifespan that matches its host if cognitive function is to be maintained. An insufficient mtDNA population would lead to an energy deficit within the cell, thus, mtDNA must replicate in order to mitigate natural attrition. However, like any replicon, the mitochondrial DNA wild-type (mtDNA_*wild*_) is subject to mutation giving rise to variant species (mtDNA_*del*_). As mtDNA_*del*_ are also replicons they will compete with mtDNA_*wild*_.

Indeed, experimental evidence suggests mtDNA exhibit *selfish* behaviour [7, 15, 37] where the environment imposes competition for limited resources. In this paper, we model the mitochondria as a simple organelle of fixed mtDNA capacity. In our simulation, we set the capacity such that it can easily accommodate a copy number sufficient to maintain necessary energy levels for the cell. mtDNA_*wild*_ maintains this copy number through an energy deficit/surplus feedback system. However, mtDNA_*del*_ proliferate selfishly which can lead to the consumption of the excess capacity. At this point, competition for space ensues between mtDNA_*wild*_ and mtDNA_*del*_ that is Darwinian in nature [16]. The ultimate survival of the cell depends upon the survival of the mtDNA_*wild*_ where small advantages can tip the competition outcome in favour of mtDNA_*del*_, even if it is to the detriment of the mtDNA_*del*_ population itself.

Proliferation of mtDNA_*del*_ has been observed in post-mitotic cells [28], however, despite an overall increase in the total mtDNA, an amount of mtDNA_*wild*_ is maintained at a steady level to fulfill ATP requirements. By necessity there must be a feedback mechanism by which the organelle can detect and respond to a deficit of ATP by increasing the amount of mtDNA [1].

In this paper we use simulation methods to study mitochondrial dysfunction caused by the proliferation of mtDNA_*del*_. The simulation software is based upon our previous publication [21] and has since been adapted to simulate a number of mtDNA proliferation mechanisms. Our aim is to compare different mechanisms and assess their suitability in predicting current empirical observations. Specifically, we compare three mtDNA proliferation mechanisms namely, random drift (RD), replicative advantage (RA) and vicious cycle (VC) with respect to cell loss rates and heteroplasmy.

We describe the three mechanisms below:

- RD: In this mechanism mtDNA_*del*_ are benign and have no replicative advantage. mtDNA_*del*_ have no adverse effect on the organelle other than to take up space. However, when the capacity of the organelle is reached, the ability of mtDNA to clone is disrupted. Despite the smaller size of mtDNA_*del*_ we keep replication times constant across *all* species of mtDNA. As mtDNA_*del*_ have no replicative advantage proliferation is *purely* a result of random drift.
- RA: mtDNA_*del*_ are benign but have a replicative advantage. mtDNA replication times are set proportional to their size (where larger deletions are smaller in size). In RA random drift is still a factor in mtDNA_*del*_ proliferation.
- VC: mtDNA_*del*_ are pathogenic, in that, mtDNA_*del*_ proliferation effects the mutation rate over the cell’s life time, whereas, with RD and RA the mutation rate is kept constant. With VC the mutation rate increases with the number of mtDNA_*del*_. This in turns gives rise to the *vicious cycle*. Random drift and replicative advantage still apply within VC.

For each mechanism described above we run the simulation over a range of mutation probability parameters and observe the effect on mtDNA_*del*_ proliferation within an organelle. We also observe the cell’s longevity, that is, a cell survives for the entire lifespan of its host or it dies prematurely due to being overrun by mtDNA_*del*_. From this we determine the cell loss rates associated with mutation rates. There is little consensus in the literature as to what cell loss rates, both global and local, constitute dementia [2, 11, 13, 43]. For the purpose of this paper we select a target of 20-40%, where 20% is the onset of dementia and 40% is advanced. Below 20% is merely healthy aging and above 40% it is likely the host would not have survived.

We also examine heteroplasmy. Historically, mutant proliferation was attributed to a unique clonally expanded mutant [25] which is what had previously been observed. This led to VC being discounted as viable proliferation mechanism as it predicts high levels of heteroplasmy. However, we argue that any proliferation mechanism that relies on random mutations would yield some level of heteroplasmy. Indeed, more recent studies of dementia patients has revealed that the population of mtDNA_*del*_ is *not* a single clone [6, 9]. Furthermore, ultra-deep mapping of the mitochondrial genome in neuronal cells indicates that neurons contain a heterogeneous pool of low-frequency mtDNA_*del*_ with 1–4 abundant species [31, 34].

Thus our inputs to the system are mutation probability parameters and our outputs are cell loss rates. However, we do not compare proliferation mechanisms for a common set of input parameters, rather we select a range a parameters for each mechanism that yields a *common* output. That is, we select parameters for each mechanism that result in cell loss rates that yield a range of cognitive dysfunction from healthy aging to advanced dementia.

The contribution of this paper to the topic of mtDNA_*del*_ proliferation and the onset of dementia is three-fold:

- Creation of a simulated environment where mtDNA expire, replicate, mutate and compete. Within this environment, mtDNA_*del*_ behave *selfishly* thus forming a Darwinian “microcosm” [16].
- Determination of cell loss rates for three mtDNA_*del*_ proliferation mechanisms, namely, RD, RA, and VC for a range of mutation rates.
- Comparison of each mechanism with regard to heteroplasmy. We evaluate the results against current empirical observations in order to assess which mechanism best predicts mtDNA_*del*_ proliferation and its effect on cognitive dysfunction in a human host.

Results data and source code for the simulation is available on Github: git@github.com:agholt/closs.git.

## 2 Simulator

In this paper we simulate the mtDNA population within a pseudo organelle based on the mitochondrion. Our simulator is adapted from the one previously developed [21] to implement RD and RA mechanisms which we further developed to implement VC. For each mechanism we observed the proliferation of mtDNA_*del*_ over time against a population of mtDNA_*wild*_. The dynamics of the system are governed by four principles:

- Expiration. mtDNA are relatively short lived compared to cells (particularly post-mitotic) with a lifespan in the order of 10-30 days. In our simulation, mtDNA expire according to a specified half-life of 30 days.
- Replication. Cells need to maintain a population of mtDNA despite their shorter lifespan. mtDNA replicate by cloning, thus, replenishing the population within the organelle.
- Mutation. When mtDNA undergo cloning, mutation can occur which gives rise to a new species of mtDNA. In this paper we focus solely on mtDNA_*del*_. mtDNA_*del*_ expire and replicate in the same way as mtDNA_*wild*_, thus, populations of mutant species can grow in the organelle alongside the mtDNA_*wild*_. mtDNA_*del*_ are also subject to further mutation yielding larger deletions.
- Competition. We place a limit on the size of the mtDNA population, within the organelle. When the organelle reaches capacity, cloning is disabled. Space within the organelle is freed up only when some mtDNA have expired. This creates competition between the species for *space*. If mtDNA_*wild*_ are overwhelmed by mtDNA_*del*_ the cell dies as its depleted mtDNA_*wild*_ is unable to generate sufficient energy.

The simulator operates in discrete time intervals *t*, where *t* is 15 minutes. Each simulation is run for 100 years. At time interval *t* = 0 the simulator is initialised with a population of *N*_*wild*_ = 2, 000. This mtDNA_*wild*_ copy number is maintained via an ATP feedback mechanism such that the mtDNA_*wild*_ population produces the requisite amount of energy for the cell.

In each subsequent interval *t >* 0, mtDNA can replicate and randomly mutate. Random deletion mutations lead to heteroplasmy within the organelle such that the overall mtDNA population comprises mtDNA_*wild*_ and mtDNA_*del*_. Deletion mutations in our simulation are idealised in that an mtDNA_*del*_ is half the size of its parent.

The relatively short lifespan of mtDNA compared to that of the host necessitates replication such that expired mtDNA_*wild*_ are replaced to maintain ATP levels. Consequently, mtDNA_*del*_ also replicate which leads to their proliferaton.

The limit imposed on the population size in the organelle will lead to competition for space between the different species. One or more species of mtDNA_*del*_ can proliferate and overwhelm mtDNA_*wild*_ leading to the eventual death of the cell as the required ATP level can not be maintained. A more detailed description of the simulation model is given in the subsections below.

### 2.1 Expiration

mtDNAs expire over time due to *aging*. mtDNA aging is simulated by assigning a time-to-live (TTL) to each mtDNA which is decremented each time interval *t* according to a Bernoulli trial success:

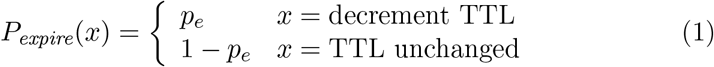

When the TTL reaches zero, the mtDNA expires. The TTL and the Bernoulli trial probability *p*_*e*_ are calibrated to yield a half-life of 30 days.

### 2.2 Replication

In each time interval *t >* 0, all mtDNA can replicate if *all* of following conditions are met:

1. Organelle-wide cloning is enabled by the ATP deficit/surplus feedback mechanism. The host cell has ATP requirements, a deficit in ATP activates cloning as a means of increasing ATP production. Conversely, cloning is deactivated when there is a surplus of energy. This condition affects the ability of all mtDNA in the organelle to replicate.
2. The initial mtDNA population is below the capacity of the organelle. If *C*_*max*_ is the capacity of the organelle, then cloning is enabled only if |*mtDNA*|*t* ≤ *C*_*max*_, where |*mtDNA*|*t* is the number of mtDNA at any given interval *t*. If the mtDNA population exceeds *C*_*max*_ cloning is disabled organelle-wide. Cloning is re-enabled once the mtDNA population drops below *C*_*max*_ due to mtDNA expiry.
3. The mtDNA is not already undergoing replication. The cloning process takes a finite amount of time to complete and an mtDNA is not eligible to replicate until the end of the cloning process. This condition is applied per-mtDNA. Replication time depends upon the mechanism (RD, RA or VC) being simulated and is discussed in more detail below.
4. Bernoulli trial success for a given cloning probability. For each time interval *t* and mtDNA, a Bernoulli trial is run. A success enables the mtDNA to undergo replication:

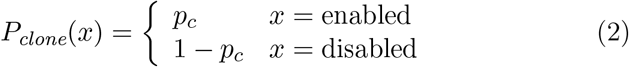

where *p*_*c*_ = 0.01. This condition is applied per-mtDNA.

The cell’s demand for energy, in the form of ATP, is modeled as an external “consumer” (to the organelle) that periodically requests a quantity of ATP. Every time interval *t* each *w* ∈ mtDNA_*wild*_(*t*), “produces” an amount of ATP *E*_*w*_ which is added to a common ATP pool for the cell, *E*_*ATP*_(*t*). The consumer extracts ATP *E*_*cell*_ from the pool in order to fulfil the cell’s energy requirements:

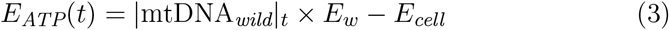

If ATP for the cell is in deficit (*E*_*ATP*_(*t*) *<* 0), cloning is enabled but if there is a surplus (*E*_*ATP*_(*t*) ≥ 0) then cloning is disabled. Thus we use this ATP deficit/surplus feature to maintain the wild-type copy number, which in our paper is 2,000 mtDNA.

In reality, we appreciate that mtDNA sequences do not produce ATP directly and there is no common “ATP pool”. Nor does the external consumer exist as an single biological entity. Our model is merely a convenient abstraction of the energy usage process within the cell.

Replication of an mtDNA takes time and an mtDNA undergoing cloning is placed in a *busy* state so that it cannot re-enter cloning until the busy period has elapsed. Replication times depend upon the model we are simulating. For RA and VC the replication time is proportional to the mtDNA’s size. An mtDNA_*wild*_ is 16,569 nucleotides in size and takes two hours to replicate. Any mutated child is half the size of it’s parent and, therefore, takes half the time to replicate.

For any *m* ∈ mtDNA, we compute the replication time (busy period) *T*_*clone*_(*m*) with the expression:

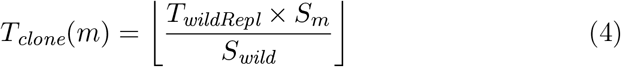

where *S*_*m*_ is the size (in nucleotides) of *m*, S_*wild*_ is the size of mtDNA_*wild*_ (16,569) and *T*_*wildRepl*_ = 8 time intervals (two hours).

For RD the replication time is kept constant for all mtDNA regardless of size. That is, both mtDNA_*wild*_ and mtDNA_*del*_ take two hours to replicate.

### 2.3 Mutation

A replicated mtDNA is either an exact clone of its parent or a deletion mutation. As mutants can themselves mutate, the parent may be a mtDNA_*del*_ of some other mtDNA_*del*_. Mutation takes place upon the success of a Bernoulli trial of probability *P*_*mutate*_:

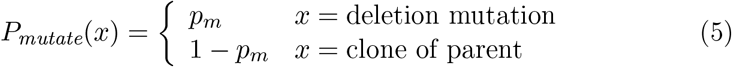

The value of *p*_*m*_ (at any time *t*) is given by the expression:

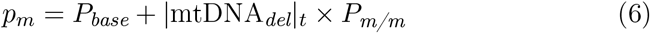

where *P*_*base*_ is the base mutation probability (per hour) and *P*_*m/m*_ is the per-mutant mutation probability (per hour). |mtDNA_*del*_|_*t*_ is the number of mtDNA_*del*_ at time interval *t*. For RD and RA *P*_*m/m*_ = 0. For VC *P*_*m/m*_ is a non-zero value, thus, as the number of mtDNA_*del*_ increases, *p*_*m*_ increases, thereby causing a vicious cycle.

### 2.4 Competition

Enforcing a maximum capacity (*C*_*max*_) in the organelle creates competition for space between species. Initially there is an abundance of space within the organelle and there is spare capacity for all species of mtDNA. However, as the mtDNA_*del*_ population grows, it eventually reaches *C*_*max*_. We define the moment where the population reaches *C*_*max*_ for the first time as the *competition-point*. At this point, space within the organelle is scarce and each species of mtDNA must compete for it. When the organelle enters competition, selfish proliferation of mtDNA_*del*_ impacts the ability of mtDNA_*wild*_ to maintain energy levels within the cell. The cell dies if the mtDNA_*wild*_ population drops below a threshold (in our simulation, *N*_*wild*_*/*3).

### 2.5 Model Parameters

Table 1 shows the *fixed* parameters that are used in every simulation run. Table 2 shows the parameters that varied across simulations, namely, *P*_*base*_ and *P*_*m/m*_ which are the terms in Eq (6). Note that, our aim is not to compare “probability” parameter values across mechanisms. Indeed, for VC we vary an entirely different parameter, namely the per-mutation mutation probability (*P*_*m/m*_), than for RD and RA where we vary *P*_*base*_.

**Table 1:**
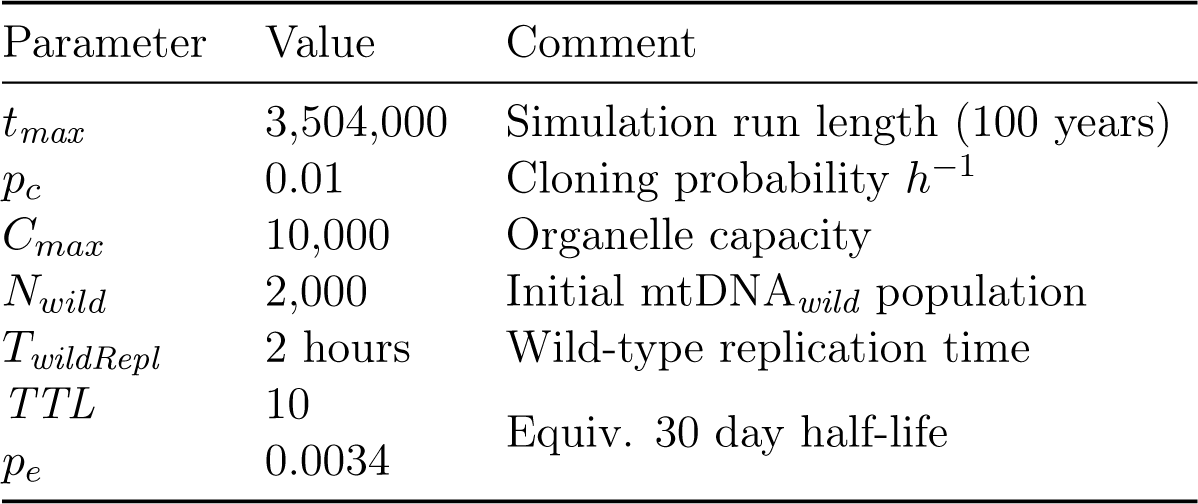
Simulation parameters, fixed for all simulations.

**Table 2:** Mutation probability parameters. For RD and RA *P*_*base*_ is varied and *P*_*m/m*_ is kept constant. For VC *P*_*m/m*_ is varied and *P*_*base*_ is kept constant.

| Mech | $P_{base} (h^{-1})$ | $P_{m/m} (h^{-1})$ |
| --- | --- | --- |
| RD | $(2, 3, 4, 5, 6) \times 10^{-3}$ | 0.0 |
| RA | $(2, 4, 6, 8, 10) \times 10^{-4}$ | 0.0 |
| VC | $10^{-4}$ | $(2, 4, 6, 8, 10, 12) \times 10^{-6}$ |

Our aim in this paper is to select the parameter values for each mechanism that yield typical levels of cognitive dysfunction (in terms of cell loss rates) experienced by a human host living 100 years. From this, we can compare the rate of decline for each mechanism and levels the of heteroplasmy that result. We then make an assessment as to which mechanism is the most suitable for predicting human cognitive dysfunction.

## 3 Analysis

In this section we present the results of cell death rates for each of the proliferation mechanisms. The simulations were conducted over a range of values of *P*_*base*_ and *P*_*m/m*_. For RD and RA we set *P*_*m/m*_ = 0 *h*^-1^ and varied *P*_*base*_, whereas for VC we set *P*_*base*_ = 10^−4^ *h*^-1^ and varied *P*_*m/m*_. The values of *P*_*base*_ and *P*_*m/m*_ are summarised in Table 2.

For each mechanism and each associated range of values, *P*_*base*_ and *P*_*m/m*_, the simulation was run 100 times and the number of cell deaths was recorded. A cell is deemed to have died if the mtDNA_*wild*_ population falls below a threshold of *N*_*wild*_*/*3 before the end of the simulation run and survived, otherwise.

### 3.1 Cell Loss

Figure 1 shows the cell loss rates for RD, RA and VC. Cell loss rates were derived by taking the ratio of cells that died before the host and those that survived up to the lifespan of the host:

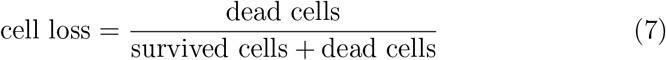

**Figure 1:**
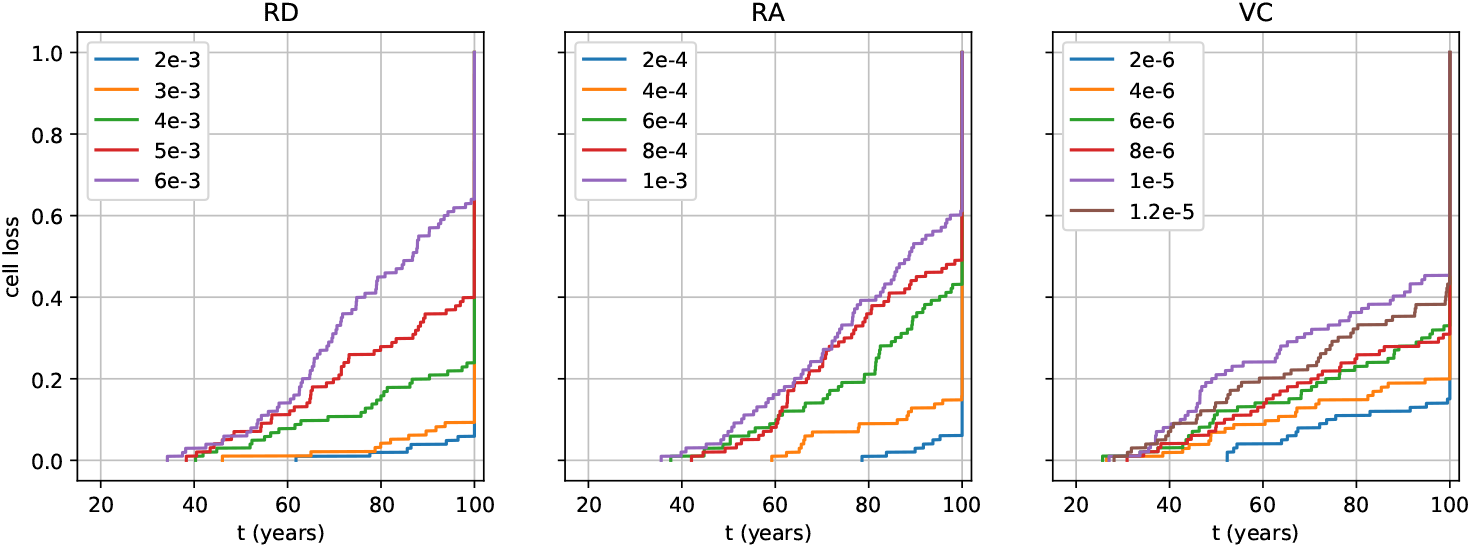
Cell loss. For RD and RA, *P*_*m/m*_ = 0 is kept constant while *P*_base_ is varied over the values (2, 3, 4, 5, 6) × 10^−3^ and (2, 4, 6, 8, 10) × 10^−4^, respectively. For VC, *P*_base_ = 10^−4^ is kept constant and varies over the values *P*_*m/m*_ = (2, 4, 6, 8, 10, 12) × 10^−6^.

For RD, *P*_*base*_ = (2, 3) × 10^−3^ *h*^-1^ yields low levels of cell loss late on in life (*>* 80 years) and, while there may be some cognitive decline, this would be considered merely the result of healthy aging. At *P*_*base*_ = 4 × 10^−3^ *h*^-1^ early signs of dementia may be experienced but not until the host’s late-eighties. However, at *P*_*base*_ = (5, 6) × 10^−3^ *h*^-1^ dementia occurs in the host’s early sixties. Furthermore, there is a rapid decline and, as the host enters its eighties, dementia is quite severe at 40% cell loss. While the simulation runs until age 100, it is unlikely that a host with such advancing levels of cell loss would live much beyond this point.

RA shows a similar pattern to RD except at much lower values of *P*_*base*_. Dementia first arises in the host’s early eighties for *P*_*base*_ = 6 × 10^−4^ *h*^-1^. This is followed by rapid decline until the end of the simulation. For *P*_*base*_ = (8, 10) × 10^−4^ *h*^-1^ dementia begins in the host’s early sixties and gets progressively worse into the host’s eighties. Again, we would not expect the host to live as long as the simulation run.

We note, that for RD and RA, for their respective values of *P*_*base*_, the first signs of cell loss occur in the host’s early thirties. VC yields a slower decline but cell loss begins at a much earlier age compared to RD and RA. Dementia starts in the host’s seventies for *P*_*m/m*_ = (6, 8) × 10^−6^ *h*^-1^ but starts in host’s early-fifties/late-forties for *P*_*m/m*_ = (1, 1.2) × 10^−5^ *h*^-1^.

### 3.2 Heteroplasmy

In this sub-section we compare heteroplasmy of each mechanism. First we present the results for total number of mutant species over the lifetime of a cell (died or survived). We use bootstrap methods [12], that is, resampling with replacement, to generate a sampling distribution of means for the number of mutant species. The graph in Fig 2 shows a boxplot for the number of mutation species over the range of mutation probabilities for the three mechanisms. Furthermore, we divide the results into cells that died and cells that survived.

**Figure 2:**
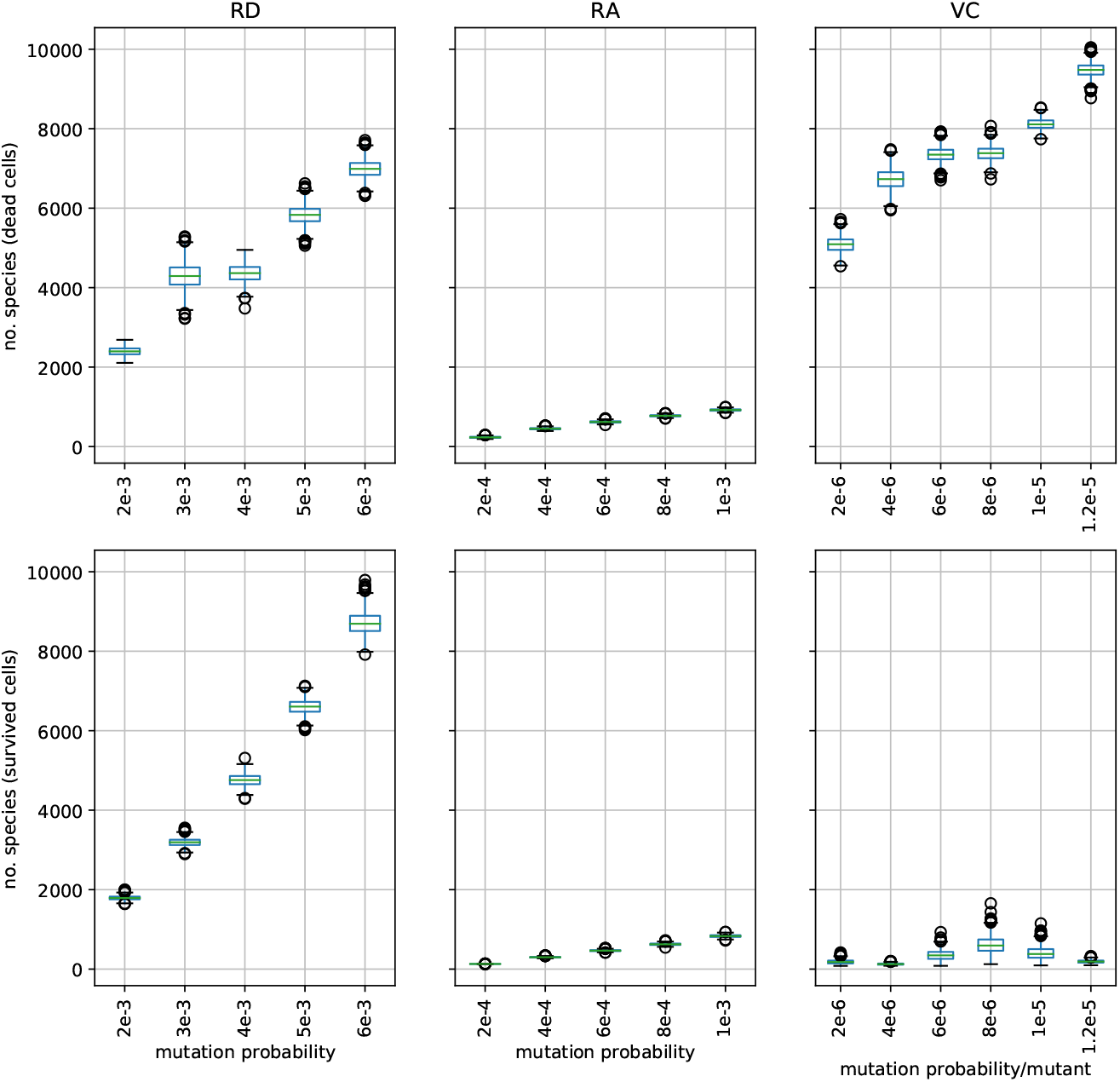
Heteroplasmy. Number of mutant species over cell lifetime (survived and died).

We can see that for RD the number of mutant species is high in both cells that died and survived, whereas for RA the number of mutant species is much lower (for both died and survived). For VC, mutant species are high for cells that died but low for cells that survived.

Clearly RA, even for cells that die, yield low levels of heteroplasmy. We attribute this to the lower mutation probabilites of RA to drive mutation proliferation within the cell. RD requires much higher mutation probabilities than RA to achieve comparable cell loss and high mutation probability leads to increased heteroplasmy. Similarly, with VC, when the system is pushed into vicious cycle, the mutation probability increases, which also gives rise to higher heteroplasmy. This pattern is predominant in dead cells. In contrast, cells that survived have very low levels of mutation (rarely do they reach the point of competition) and, consequently, heteroplasmy is low.

High levels of heteroplasmy are an inevitable consequence of high mutation rates. High mutation rates cause high cell loss congruent to severe cognitive dysfunction for both RD and VC. RA can, however, cause this level of cell loss at much lower mutation rates which results in lower heteroplasmy.

Finally, we examine the state of heteroplasmy of cells that have died at the time of their death (when the simulation ends due to mtDNA_*wild*_ dropping below the *N*_wild_*/*3 threshold). Cells that have died have high levels of mutation (as they have reached the point of competition and, eventually, the wild-type is overwhelmed). As well as the number of mutant species that are currently extant at this point, we are also interested in the evenness of the distribution of mutants across the species. For the former, we show the mean number of mutant species in existence just prior to the cells death; and for the latter, we use information entropy [8], where entropy is given by:

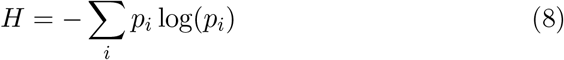

and *p*_*i*_ is the normalised number of mutants in the *i*^*th*^ species:

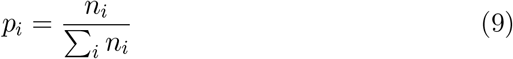

A high entropy value indicates an even distribution of mutant species whereas as a low value indicates an uneven distribution.

Figure 3 shows a boxplot for the heteroplasmy and entropy for dead cells at the time of their death. We use bootstrap methods (resampling rate = 10,000) to get a sampling distribution. Note, we only show results for mutation probabilities (*P*_*base*_ and *P*_*m/m*_) that yield the levels of cell loss that would result in cognitive dysfunction (as previously defined). For low mutation probabilities, cell deaths are so infrequent that it is difficult to derive meaningful statistics (even with re-sampling techniques).

**Figure 3:**
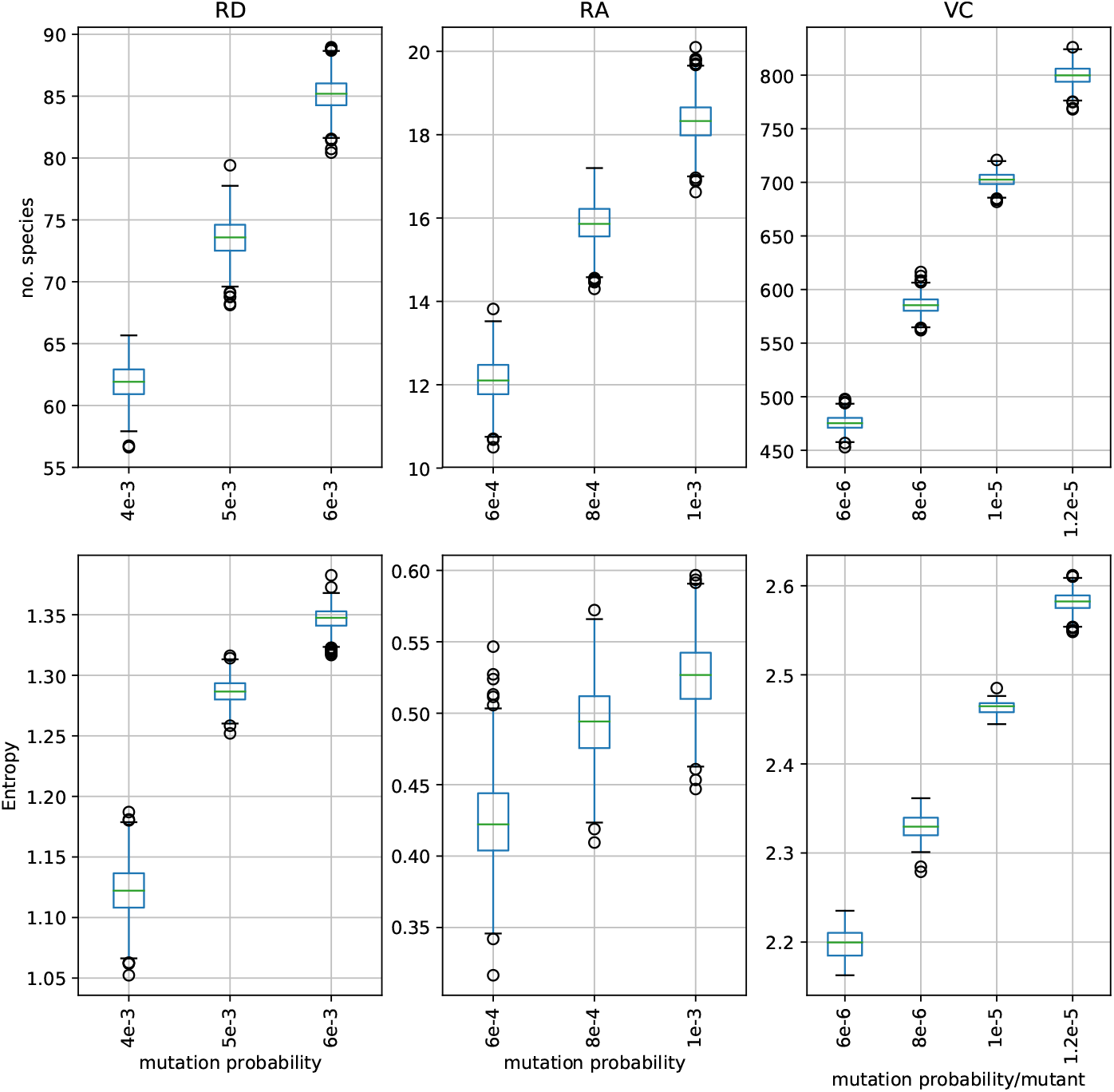
Heteroplasmy and entropy of dead cells at the time of death.

From the graph in Fig 3, we can see the RA heteroplasmy is relatively low. It is an order of magnitude higher for RD and another order of magnitude higher for VC. The low entropy values for RA indicate that there is an uneven spread of mtDNA_*del*_ across species. For RD and VC, which yield higher entropy values, mtDNA_*del*_ are more evenly spread.

## 4 Discussion

Our simulation operates on the basis that the mtDNA_*del*_ are a causal factor in the development of dementia. Indeed, the proliferation of mtDNA_*del*_ has been observed in the brains of people with Alzheimer’s disease [20, 36]. The age of onset and rate of progression of neurodegenerative dementias, such as Alzheimer’s, is well documented [39] and which we use to inform our expectations regarding the outcome of our simulations. For each mechanism we select mutation rates (through the parameters *P*_base_ and *P*_*m/m*_) that yield a range of cognitive dysfunction, namely, from healthy aging to severe dementia.

Our results support the observation that dementia seldom occurs below the age of 65 [36]. In addition, further evidence indicates that part of the pathogenesis of neurodegenerative dementias is that the mtDNA heteroplasmy, *in vivo*, consists of the predominance of a small number of mtDNA_*del*_ species present at a high copy number, with a low background of heteroplasmy [27, 34].

The cell loss results in Fig 1 show that RD and RA have a 20-40% cell loss over the age range of 60 to 80 years (approximately) with sufficiently high mutation rates. VC shows a slightly slower cognitive decline than RD and RA and an earlier onset of dementia at late-forties/early-fifties. Note, the straight vertical line at the right hand edge of the graphs in Fig 1 indicate the cell survived rates.

The mutation rates required to achieve increases in mtDNA_*del*_ proliferation vary considerably between the three mechanisms. RD requires a *P*_base_ value an order of magnitude greater than RA. For VC, we use a relatively low value of *P*_base_ but vary *P*_*m/m*_. The value of *P*_base_ = 10^−4^ is, likely, too low to achieve significant proliferation such that competition is enacted. However, using *P*_*m/m*_ = 10^−5^ as an example, the mutation probability (using Eq 6) is 0.02 when the mtDNA_*del*_ copy number reaches 2,000 and rises to 0.09 at 8,000. Which are greater than any *P*_base_ value used for RD or RA.

There has been much progress in understanding the mutation rate of mtDNA, for example, it is known that mtDNA mutation rates are greater than those of nuclear DNA [47]. However, most studies are over a lifetime or multi-generational [5] and present *overall* mutation rates after a lifetime of replication and selection. It is, therefore, unlikely, that long-term mutation rate values are applicable to those at replication cycle scales as used in our simulation.

Heteroplasmy is a general term for the presence of different species of the mtDNA in the cell. However, we were not just interested in the amount of mutant species, but how per-species copy numbers were distributed. For this, we employed information entropy [8], where low entropy values indicated an uneven distribution of heteroplasmy while larger values indicated increased evenness. As stated above, current research reveals a small number of clonal expansions with a low background heteroplasmy in dementia sufferers. Thus we were looking for the mechanism that resulted low entropy for heteroplasmy.

Our results show that heteroplasmy increases with mutation rate. There are distinct differences in the heteroplasmy between the three mechanisms. The most notable result is that heteroplasmy is much lower for RA than RD and VC.

We can compare heteroplasmy between cells that survived and those that died. While for RD, heteroplasmy is similar for both cells that died and survived, it is higher for survived cells than for dead. This is because survived cells live longer than cells that die before the end of the simulation, thus more mutants arise in those additional years. Heteroplasmy for RA is slightly lower for survived cells than died (but low for both).

For cells that survived in the VC simulation, heteroplasmy is low. This is because there was little proliferation of mutants as it is unlikely that the mechanism actually entered *vicious cycle*, keeping mutation rates low. If the mechanism enters vicious cycle, the mtDNA_*del*_ proliferation pushes the organelle into competition resulting in the cell’s eventual demise. The consequence of the increase in mutation rates caused by vicious cycle leads to higher heteroplasmy.

We implement a capacity limit in our simulation that, when reached, disables cloning organelle-wide even if the ATP is in deficit. Cloning can only be re-enabled after some mtDNA attrition has freed up space in the organelle. At the point of competition, ATP production is disrupted as mtDNA_*wild*_ compete with mtDNA_*del*_ for space. This usually marks the “beginning of the end” for mtDNA_*wild*_ as its copy number depletes until it drops below the *N*_wild_*/*3 threshold and the cell dies.

This implementation feature is supported by experimental evidence of selection and competition between mtDNA species for limited resources [16, 29]. In reality, the resource over which mtDNA compete may not actually be space, nevertheless it is an adequate abstraction for the purpose of our simulation. Furthermore, the modular nature of our model makes it amenable to simulate different selective conditions and explore various hypotheses.

## 5 Conclusions

In this paper we investigated cell loss due to mtDNA_*del*_ proliferation for three mechanisms, namely, random drift (RD), replicative advantage (RA) and vicious cycle (VC).

Mutation rates (via mutation probability parameters) were adjusted to achieve cells loss rates congruent with dementia, that is, 20-40%. All three mechanisms predicted that dementia was a disease of old age. Thus, we propose that a human host that is predisposed to high mutation rates is similarly disposed to dementia in advanced age.

That larger deletions have a replicative advantage has been disputed and the reason given is the scale difference between replication time and half-life of mtDNA [24]. We demonstrated in a previous paper that mtDNA_*del*_ *do* have a replicative advantage despite replication time being of the order of a few hours and half-life in tens of days [21]. The results of this research also show strong evidence for replicative advantage given that there is comparable proliferation between RD and RA despite RA needing much lower mutation rates.

The notion of a single clonal expansion accounting for the entire proliferation of mtDNA_*del*_ has been superseded by research that reports a small number of clonal expansions with a low background of heteroplasmy [30, 32, 34, 46]. This is consistent with the expectation that, in a biological system subject to random mutation, heteroplasmy is ubiquitous.

In RD and VC high levels of heteroplasmy were observed in cells where mtDNA_*del*_ proliferated. However, for RA, heteroplasmy was much lower for comparable proliferation. Furthermore, the entropy analysis suggests a small number of predominant mtDNA_*del*_ clones with low levels of heteroplasmy. Thus, replicative advantage would appear to be the most suitable mechanism out of the three for predicting mtDNA_*del*_ proliferation within a human host.

